# Fractional re-distribution among cell motility states during ageing

**DOI:** 10.1101/2020.04.29.069286

**Authors:** Jude M. Phillip, Nahuel Zamponi, Madonna P. Phillip, Jena Daya, Shaun McGovern, Wadsworth Williams, Katherine Tschudi, Hasini Jayatilaka, Pei-Hsun Wu, Jeremy Walston, Denis Wirtz

**Affiliations:** Department of Chemical and Biomolecular Engineering, Institute for NanoBiotechnology (INBT), and Johns Hopkins Physical Sciences Oncology Center, Johns Hopkins University, Maryland, USA; Department of Medicine, Hematology and Oncology Division, Weill Cornell Medicine, New York, USA; Department of Medicine, Division of Geriatric Medicine and Gerontology, Johns Hopkins University School of Medicine, Maryland, USA; Departments of Oncology and Pathology, Sidney Kimmel Cancer Center, Johns Hopkins University School of Medicine, Maryland, USA; Departments of Biomedical Engineering, Johns Hopkins University, Maryland, USA

## Abstract

Ageing in humans is associated with a decreased capacity to regulate cell physiology. Cellular properties, such as cell morphology and cell mechanics, encode ageing information and as a result can be used as robust ageing biomarkers. Using a panel of dermal fibroblasts derived from healthy donors spanning a wide age range, we observe an age-related reduction in average cell motility, which we show is not due to the decreased motility of all cells, but results from fractional re-distribution among motility states. By taking advantage of the single-cell nature of our motility data, we show that cells can be classified based on spatial and activity patterns that define age-dependent motility states. These findings highlight an important feature of ageing cells shown by the decrease in the heterogeneity of cell movement in older adults, that potentially offer new mechanistic insights into the ageing process and avenues for novel biomarker development.

## INTRODUCTION

Ageing can be defined as the accumulation of damage with the passage of time that limits the ability of organisms, organs and tissues to absorb and rebound after perturbations and stressors^1–3^. In humans, normal ageing is associated with diverse physiological changes that influence the magnitude and rates of progressive decline among individuals. These include the decreased abundance and activity of circulating cytotoxic immune cells, slower gait speed and declines in cardiorespiratory fitness^4–8^. There is also a high degree of variability between individuals, which suggests that there is not a uniform aging phenotype. A growing body of evidence shows that the interactions of intrinsic and extrinsic factors such as molecular states^9–12^ (e.g. epigenomic) together with environmental determinants and macroscopic stressors^13–15^ (e.g. disparities and socio-economic factors) contribute to the ageing processes in individuals^16^. However, it remains unclear how the underlying molecular states of an individual relate to their clinical outlook at the organ/tissue level in the context of ageing. We postulate that studying age-associated changes at the intermediate length scale of cells themselves – between the larger length scale of organs and tissues and the smaller length scales of molecules – may provide the missing link to understand the inter-relation of these ageing scales^3,17^.

As integrators of molecular signals, cells offer a sensitive meso-scale view of ageing, with cellular dysfunctions likely occurring prior to the manifestation of age-related disorders and diseases at the clinical level. Populations of cells typically display dynamic and heterogeneous phenotypic characteristics in the context of both health and disease^18,19^. We now know that functional biophysical properties of cells, such as cytoplasmic stiffness, force generation and morphology, which typically capture time-independent (i.e. snapshot) cellular phenotypes, encode essential ageing information that can be used as robust biomarkers of ageing in healthy individuals^17^. However, it is still unclear how exactly this information is encoded, and its potential role in developing novel approaches for precision health. We believe that the tracking and analysis of dynamic ageing phenotypes at the cellular level can provide a unique perspective that offers novel mechanistic insights into the ageing process^20^, and may provide significantly more information than snapshots obtained from fixed cells.

In this study, we analyze single-cell motility patterns of primary dermal fibroblasts obtained from healthy donors spanning a wide age range, from 2 to 92 years old. Using a combination of bulk and single-cell analysis tools, we show that cells can be classified into various “motility states” based on spatio-temporal patterns of their movements. We then demonstrate that the decrease in overall cell motility as a function of age is due to the fractional re-distribution of the abundance of cells within these motility states. Furthermore, assessing the degree of cell-to-cell variations as a function of age, we show that cellular heterogeneity is a defining feature of ageing cells, which is decreased for individuals above age 65 (older adults).

## RESULTS

### Global decrease in bulk cell motility with increasing age

Coordinated cell movements are essential for the development of tissues and organs, in homeostasis and disease^3,17^. As cells move, there is an intricate coordination of biophysical and biomolecular programs that change with age, some of which involve the modulation of cellular biomechanics, adhesion and regulated dynamics of the cytoskeleton within cells^21,22^. To elucidate possible age-related changes in cell motility patterns, we procured a panel of primary dermal fibroblasts from healthy donors spanning an age range, from 2 to 92 years old. Using time-lapse microscopy, we recorded and tracked the spontaneous movements of these cells seeded on type-I collagen-coated substrates. Analyzing cell trajectories (**Supplementary dataset 1**), we computed averaged mean-squared displacements (MSDs) of cells and found trends towards an age-dependent decrease in overall cell motility^17,23^ (**Figure 1A-B**). In particular, the values of mean-squared displacements (MSDs) evaluated at time lags of 6 and 60 min (MSD6, MSD60) and corresponding average speeds (SP6, SP60) decreased steadily with age (**Supplementary figure 1A-J**). These time lags of 6 and 60 min were chosen because they correspond to time scales shorter and longer than the average persistence time of cell motility across all ages (**Supplementary Figure 1A-D**).

**Figure 1.**
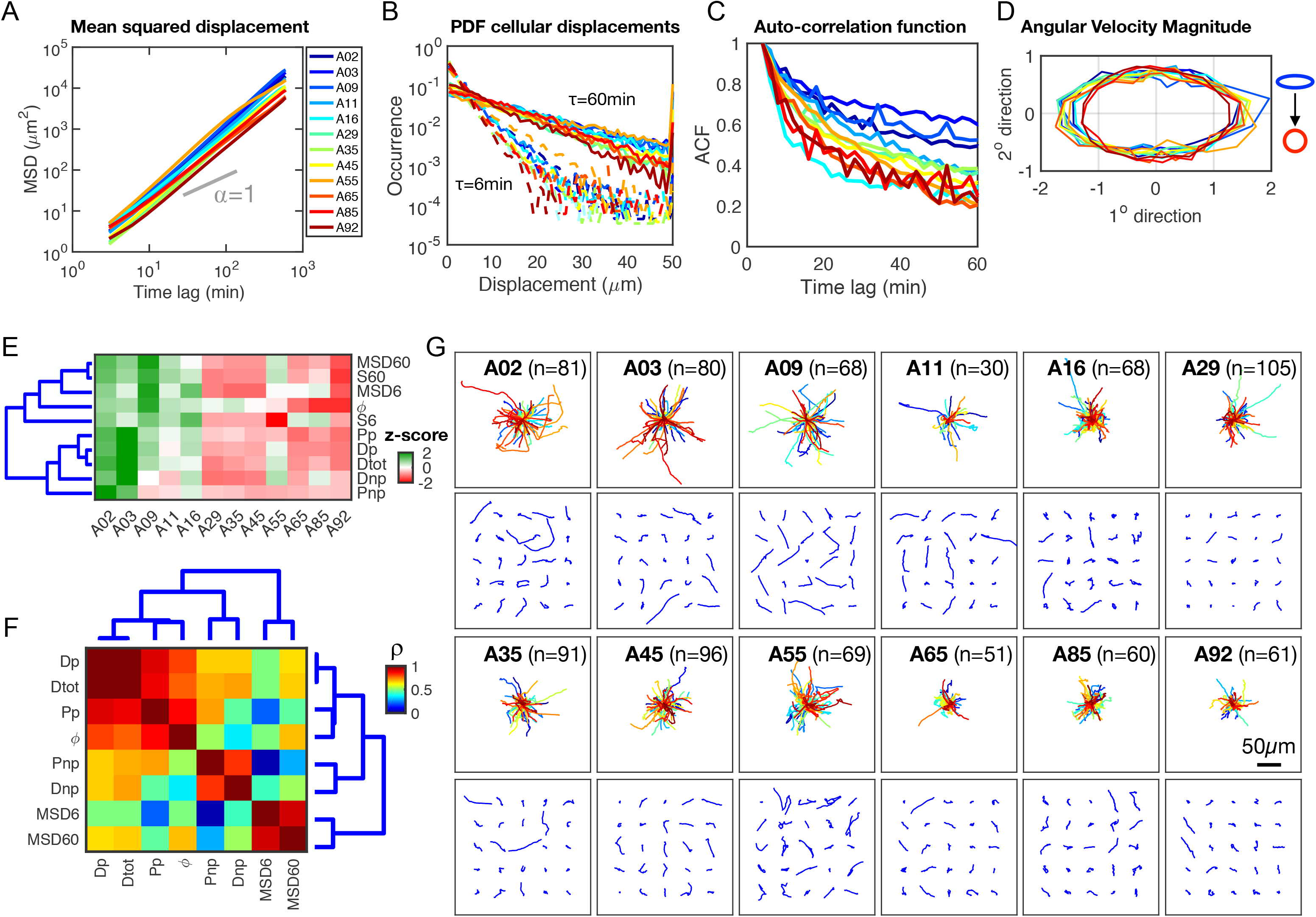
Global decrease in ensemble cell motility with increasing age. **A**. Age-dependent mean-squared displacements (MSD) of dermal fibroblasts with color code indicating the age of the donor (blue-to-red). An exponent alpha of 1 denotes pure diffusion. **B**. Age-dependent probability functions of cell displacements at time lags equal to 6 min (dashed lines) and 60 min (solid lines). **C**. Auto-correlation function of velocities (ACF) measured at 3 min time lags. **D**. Angular velocity magnitudes as a function of the age of the donor, ‘circular’ denotes similar likelihood for all angles and ‘ellipsoidal’ indicating a polarization of cell movements along primary axis of migration. Color coding in panel B also applies to panels C-D. **E**. Heatmap showing the magnitude of the correlation between motility parameters (z-score normalized) and age. Dendrogram branches indicate hierarchical clustering with ward linkages of the Cityblock distances of parameters. Red-to-green signifies low-to-high Pearson correlation coefficient. These parameters include total diffusion (Dtot), diffusion along the primary and secondary axes of cell movements (Dp and Dnp), measure of the spatial persistence—anisotropy of the cell movements (ϕ), persistence-time of cell motion along the primary and secondary axes of cell movements (Pp and Pnp), average cell speed at time lags of 6 min and 60 min (S6 and S60), and the MSD measured at time lags of 6 and 60 min (MSD6 and MSD60) (see definitions in Glossary). **F**. Heatmap showing the magnitude of correlations of motility parameters across all cells at all ages. Unsupervised hierarchical clustering highlights three key modules based primarily on the cell and displacement, range of Pearson correlation coefficients=0.39-0.94. **G**. Cell migration patterns and analysis for primary dermal fibroblasts collected from healthy donors with ages spanning 2-92 years. Top panels show origin-centered trajectories for all cells per age; bottom panels show a grid of x-y trajectories for 25 randomly selected cells per age. The number of tracked cells per sample is indicated in the upper left corner of the plot.

Building on these findings, we asked whether cells taken from young donors displayed distinct spatio-temporal motility patterns compared to cells derived from older adults. Analyzing the motility data based on the recently introduced anisotropic persistent random walk (APRW)^24,25^, a framework that recognizes that cell trajectories do not always follow random walks even on flat substrates, we first assessed the similarity of cell movements per unit time, given by the magnitude of the autocorrelation of cell velocities (details in Materials and Methods). We observed a faster decay in the autocorrelation function of successive migratory steps with increasing age (**Figure 1C**), which suggests shorter persistence times, or more frequent changes in the direction and velocity of cells with increasing age. We then asked whether this bulk decrease in motility was also accompanied by a bias in the spatial polarity of cell movements, or a similar likelihood of movements in all directions. Quantifying the angular velocity profiles of cells, we found that cells from young donors exhibited an ellipsoidal profile of angular velocities and a tendency towards a circular profile for cells from older adults (**Figure 1D**). This indicates a loss in spatial persistence and directionality of cell trajectories with increasing age. Together, these results indicate that dermal fibroblasts show a loss in both temporal and spatial persistence with increasing age, with cells from older adults moving less, with more frequent changes in their movement direction.

To systematically define bulk age-dependent motility patterns, we computed the age-associated Pearson correlation coefficients for ten parameters that describe spatio-temporal cell movement patterns as a function of age (see Materials and Methods and Glossary). These parameters include the magnitudes of cellular displacements and speeds (MSD6, MSD60, SP6, and SP60), the total diffusivity and diffusivities along primary and secondary axes of migration (Dtot, Dp, Dnp), the persistence times along the primary and secondary axes of migration (Pp, Pnp), and the spatial persistence/anisotropy (ϕ)(see Glossary for definitions). This analysis showed negative correlations between all motility parameters and age (**Figure 1E-F**, **Supplementary Figure 1K-L**), which confirms our observation of an overall decrease in cell motility with increasing age (**Supplementary Figure 1A-J**).

Given the significant age-associated changes in cell motility, we asked whether the motility patterns of individual cells could provide new insights that are not fully appreciated from the above bulk quantification. Plotting the x-y trajectories for all cells on the same length scale, we qualitatively observed the aforementioned global decrease in cell displacements, based on the footprint of cell trajectories with increasing age (**Figure 1G** top panels). However, after closely examining the movement patterns of individual cells, we observed extensive cell-to-cell variations and the presence of cells having both high net-displacement and low net-displacement patterns from the same donor (**Figure 1G** bottom rows).

### Global decrease in cell motility with age corresponds to a re-distribution among spatial clusters

Prompted by the magnitude of the observed cell-to-cell variations, we hypothesized that the age-dependent decreases in cell motility was not due to decreases in the movement of all cells, but a re-distribution of the proportions of motile and non-motile cells with increasing age. Pooling cell trajectories across all ages, we first log-normalized the motility parameters defined above and computed the z-scores per parameter to allow for comparison across the same numerical scale (**Supplementary Figure 2** and Methods). Using unsupervised hierarchical clustering (see Materials and Methods), we determined inherent cell-based and parameter-based groupings using the ‘City block’ distances along the axis of maximum variation (ward linkages). We identified three clusters based on the similarity of trends among the motility parameters for all ages (**Supplementary Figure 3A-C**), and eight spatial clusters (Pn) based on groupings of cells having similar magnitudes of displacements, diffusivity, and spatio-temporal persistence (**Figure 2A**).

**Figure 2.**
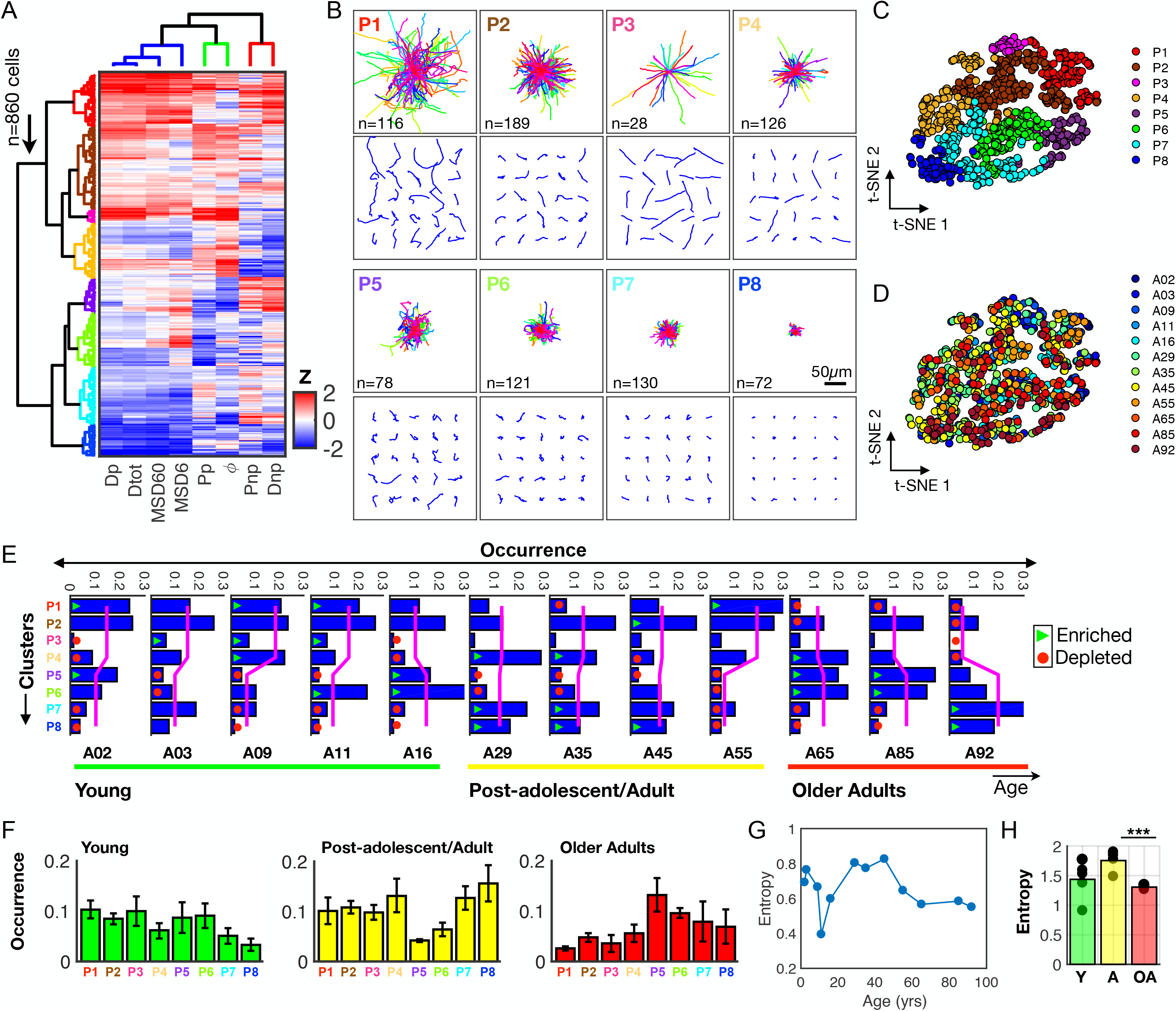
Distinct age-associated motility patterns at the single-cell level. **A**. Heatmap showing eight motility parameters (defined in the text) measured at the single-cell level across all ages (n=860 cells); each row represents a single cell and each column a motility parameter. Dendrogram branches represent hierarchical clustering using the city block distances with ward linkage along axes of maximum variation. Data shows the delineation of single cells into eight spatial motility clusters, with the column-clusters (n=3) describing the degree of cellular displacements, and spatio-temporal persistence. **B**. x-y trajectories based on the eight clusters that delineate patterns of cellular movements. Top panels show the origin-centered trajectories for all cells within clusters; bottom panels show x-y trajectories for 25 randomly selected cells per cluster, with the number of cells per cluster indicated in each plot, scale bar=50μm. **C**. Two-dimensional t-SNE visualization for the eight color-coded spatial clusters. **D**. Age-painted t-SNE plot showing the distribution of ages per spatial motility cluster. **E**. Frequency distributions indicating the fractional composition of cells per cluster as a function of age. Magenta trend lines represent the average fraction per primary branch of the dendrogram tree. The first bifurcation delineates four clusters each. Trend lines show a progressive transition in the abundance of cells across primary branches with increasing age, green triangles denote significantly enriched and red circles denote significantly depleted (p-value<0.05). **F**. Frequency distributions showing the average fractional abundances of cells per cluster separated into three groups; young, post-adolescent/adult, and older adults. **G**. Magnitude of the Shannon entropy for each individual sample with increasing age. **H**. Average Shannon entropy based on these three age groups, showing a significant decrease in heterogeneity of motility from post-adolescent/adults to older adults.

To understand the differences among these clusters and decipher what each cluster represented, we plotted the trajectories of individual cells within each cluster. Visual inspection indicated distinct patterns of cell movements in each cluster (**Figure 2B**). For instance, cluster 3 (P3) corresponded to cells having a high degree of persistence and diffusivity, whereas cluster 8 (P8) comprised cells with a non-motile phenotype characterized by small displacements, low diffusivity and low persistence (**Supplementary Figure 4A**).

Continuing to address our hypothesis of fractional re-distribution of cells, we plotted two-dimensional t-stochastic neighbor embedding (t-SNE) maps for all cells (each dot represents one cell). Coloring them based on the eight spatial clusters determined using the hierarchical clustering, we observed a confirmation of segregated groups of cells (**Figure 2C**). Using this same t-SNE layout, we then painted each cell according to their respective ages to determine whether certain clusters were defined by cells from a particular age. Interestingly, we found that cells from the same donor were intermittently distributed among all eight clusters (**Figure 2D**). To appreciate the fractional abundances of cells within each spatial cluster (Pn), we plotted the age-dependent frequency distributions for all cells (**Figure 2E**, **Supplementary table S2**), which revealed progressive age-dependent changes in the abundance of cells within various clusters. Specifically, cells from young donors tended to favor motility phenotypes described by P1-P4, while cells from older adults favored phenotypes described by P5-P8 (see magenta lines, **Figure 2E**).

Together, results indicate that there is an inherent polarization of the motility patterns based on spatial clusters exhibited by cells derived from young and older adults. In addition, cells derived from post-adolescent/adults exhibited a similar likelihood in the average abundance of cells among spatial clusters. This is denoted by the flattening of the average abundance of the two major groups of clusters (magenta lines), suggesting age-associated changes in cellular heterogeneity with age.

To quantify this age-associated heterogeneity, we grouped the cells from the donors into three age groups; young (A02, A03, A09, A11, A16), post-adolescent/adults (A29, A35, A45, A55) and older adults (A65, A85, A92), and computed the Shannon entropy, *S*, for each group, defined as^26^ (**Figure 2F**):

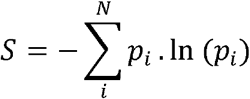

Here, *p_i_* = *n*_i_/*N* is the proportion of cells belonging to each spatial cluster Pn. The Shannon entropy measures the degree of uncertainty/disorder within a distribution. In our case the entropy is used as a surrogate for the intrinsic heterogeneity for a population of cells based on the abundance of cells within each of the defined spatial clusters. Here, the more uniform the distribution the greater the uncertainly or entropy, which is observed for cells derived from young and post-adolescent/adults (**Figure 2G-H**).

In sum, our results indicate that individual cells can be classified based on spatial patterns of their movements, with the age-associated bulk motility being approximated as the sum of weighted averages among spatial clusters per age (**Supplementary Figure 5**), and a decrease in the heterogeneity of cell movements for older adults.

### Cellular activity helps to define age-dependent cell motility

We next asked whether cells exhibited age-dependent differences in the temporal patterns of their movements. First, we defined what we call the ‘cellular activity’ for each cell in efforts to better capture the intrinsic bursty dynamics of the temporal sequences^27^, which provides insight into the fraction of time cells spent in motion or at rest. To determine the activity of individual cells, we converted each two-dimensional x-y trajectory (**Figure 3A**) into a one-dimensional displacement profile (**Figure 3B**, see Materials and Methods). We then computed the activity profile of each cell by first normalizing each one-dimensional temporal patterns to determine the magnitude (size of peaks) and frequency (number of peaks) of cellular movements. Second, we assessed whether cells displayed age-associated differences in their activity profiles that defined their movements. Pooling the activity profiles for all cells across all ages, we utilized unsupervised hierarchical clustering based on the ‘City block’ distances along the axis of maximum variation (‘ward’ linkages) to segregate individual cells into activity clusters (ACn). This analysis resulted in four activity clusters (**Figure 3C**), with each activity cluster being defined based on the frequency and magnitude of the peaks per elapsed time (**Figure 3D**). For instance, cluster 1 (AC1) was mainly defined by long periods of consistent movements close to baseline with periodic short-duration bursts, cluster 2 (AC2) exhibited frequent bursts of varying durations, and clusters 3 and 4 (AC3, AC4) exhibited extended periods of longer-duration bursts separating mainly by when the burst occurred (AC3—bursts in the beginning, and AC4—bursts towards the end of imaging time).

**Figure 3.**
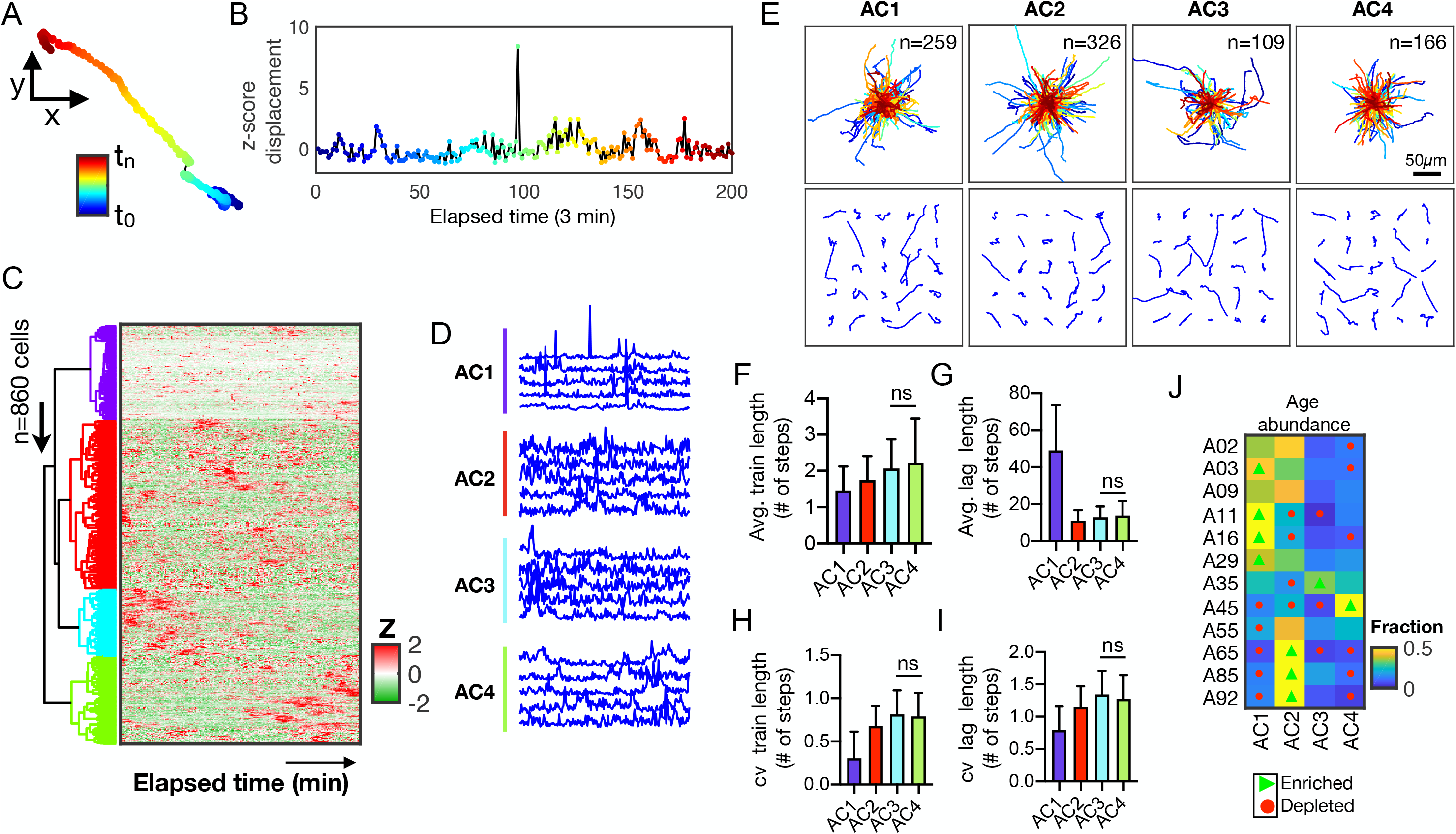
Cellular activity helps to define age-dependent motility patterns and heterogeneity. **A**. Color-coded trajectory of a representative single cell as a function of elapsed time (navy blue-to-maroon). **B**. Plot showing the corresponding one-dimensional displacement for the same individual cell, here after referred as the activity profile. **C**. Heatmap showing the cellular activity profiles for all cells, across all ages (n=860 cells). Each row represents a single cell and each column from left to right represents elapsed time. Five colored dendrogram branches represent hierarchical clustering using the city block distances with ward linkages along the axes of maximum variation. **D**. Line plots showing five representative cellular activity profiles for single cells within each activity cluster. **E**. Trajectories of cells within each activity cluster; top panel shows a grid of 25 randomly selected cell trajectories per cluster; bottom panel shows the origin-centered trajectories for all cells per cluster, with the number of cells within each cluster indicated in the upper left corner. **F-I**. Bar plots showing the extent of bursty dynamics per activity clusters; average train lengths (F), average lag lengths (G), CV of train lengths (H), CV of lag lengths (I). Error bars denote the standard deviation, with all comparisons being significantly different (p-value<0.05) except where stated with ‘ns’. **J.** Heatmap showing the abundance of cells within each activity cluster per age, green triangles denote significantly enriched and red circles denote significantly depleted (p-value<0.05).

To visually determine what these patterns represented, we plotted the x-y trajectories for cells within the five clusters (**Figure 3E**). Visual inspection of each activity cluster did not reveal distinct patterns of movement magnitudes; however, each activity cluster was populated by a mixture of both motile (i.e. high displacement) and non-motile (i.e. low displacement) cells. This suggested that both types of cells can exhibit similar activity profiles. To further qualify this observation, we took the activity profiles of each cell and asked whether applying a point-process analysis (see Materials and Methods) could better reveal a biological meaning. Taking the normalized activity profile for each cell, we set a threshold of one standard deviation above the baseline and computed the amount and frequency of movement trains (number of consecutive time steps above threshold), and the amount and frequency of the lags (number of consecutive time steps below threshold) (**Supplementary Figure 6A-B**). Compiling this binarized activity for each cell across all ages (**Supplementary Figure 6C**), we computed the distribution of trains (having a binarized activity of ‘1’) and lags (having a binarized activity of ‘0’) (**Supplementary figure 6D-E**). This analysis revealed that cells in cluster AC1 displayed significantly shorter trains and long lags (**Figure 3F-G**) compared to longer trains and short lags observed for cells in clusters AC3 and AC4. In addition, cells from AC1 were more similar in their activity based on the train- and lag-length relative to AC3 and AC4 as shown by a lower coefficient of variance (**Figure 3H-I**). Furthermore, cells from young donors were significantly enriched for AC1, with cells from older adults being significantly enriched for AC2 and significantly depleted for AC1 (**Figure 3J**, **Supplementary table S3**).

Together, the findings indicate that cellular activity can be quantified based on the magnitude and frequency of bursty dynamics in cell movements relative to their baseline. This quantitative description, together with the spatial clustering information provides a complimentary framework that describe the spatial and temporal behaviors of cells with age, with the spatial patterns describing the magnitudes of the movements, and the activity patterns describing the manner in which the cells moved.

### Cellular heterogeneity defines age-dependent cell motility states

Given the complimentary information provided by both the spatial and activity clusters, we asked whether enrichments or depletions in particular clusters could define age-dependent motility states. Compiling the frequency of cells belonging to each of the thirty-two possible motility states (based on the 8 spatial clusters and the 4 activity clusters), we plotted the frequency heatmap for each age which revealed topographic regions of high- and low-frequency states both at the single age level (**Supplementary Figure 7**), and also when categorized based on age groups (**Figure 4A**).

**Figure 4.**
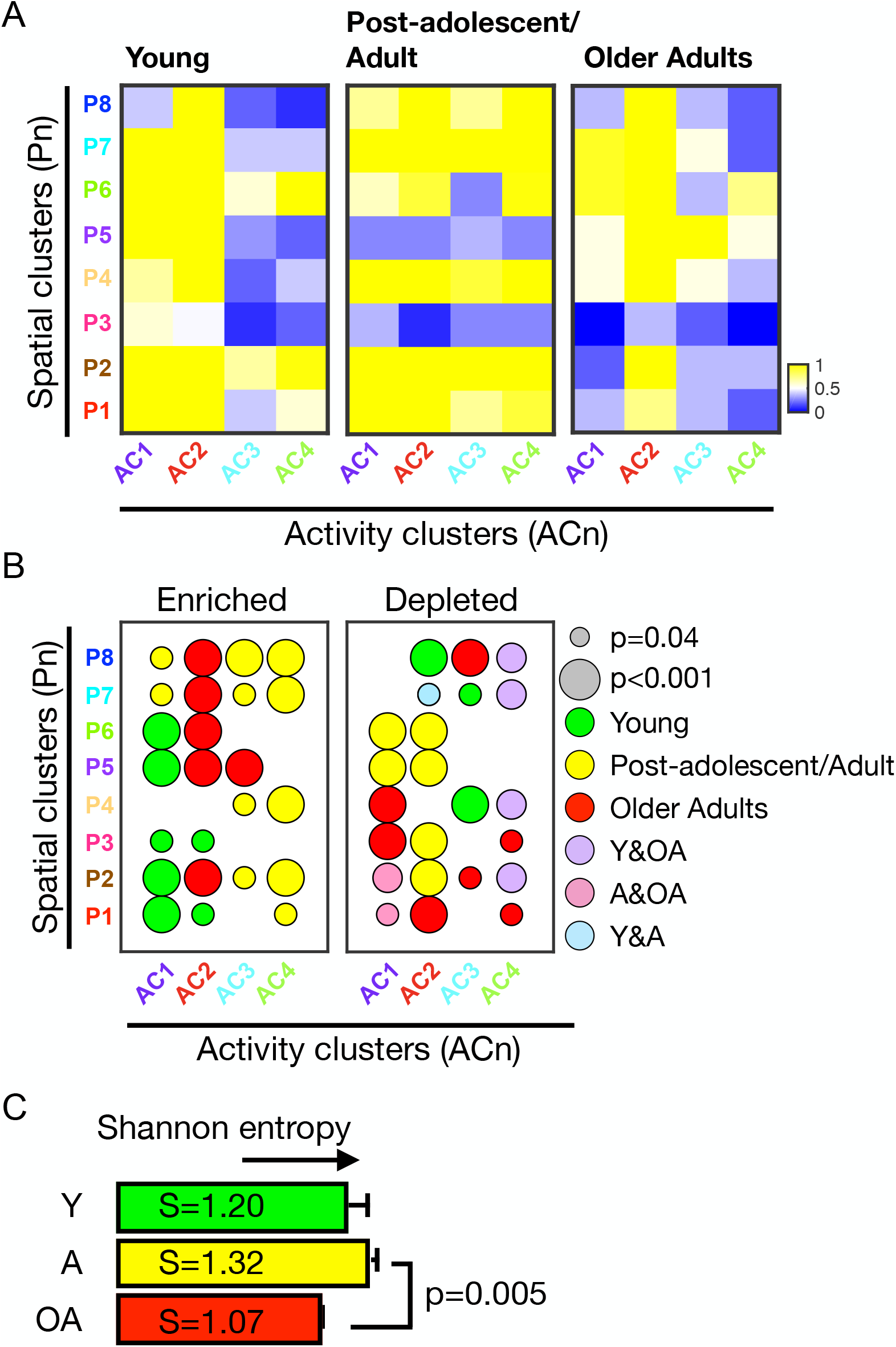
Age-dependent cell motility landscapes. **A**. Heatmaps showing the frequency per motility state defined by the eight spatial and four activity clusters: young (A02, A03, A09, A11, A16), post-adolescent/adult (A29, A35, A45, A55), and older adults (A65, A85, A92). Color scales indicate the frequencies of cells within each motility state, yellow denotes high frequencies and blue denotes low frequencies. **B**. Enrichment and depletion maps corresponding to motility states as a function of age-group. Size of the node denotes the magnitude of the p-value, and colors denote young (green), post-adolescent/adults (yellow), older adults (red). **C**. Quantification of the Shannon entropy based on age-group defined motility landscapes.

To determine whether these high- and low-frequency states were significantly enriched or depleted for cells as a function of age, we computed the statistical significance based on the null hypothesis describing the expectation at random. Utilizing a randomization test with 10,000 permutations, we asked whether the observed frequencies per state were significantly different from the expectation at random for each age group (**Supplementary Table S4-5**). These results confirmed what was visualized based on the heatmaps (**Figure 4A**), indicating significant enrichments and depletions, respectively. For instance, the dominant state observed for the co-classification of AC1 and P1, P2 and P3, and the depletion in AC3 and P4 for young donors showed p-values less than 0.05 (**Figure 4B**, **Supplementary Table S5**). Prompted by the differences in abundances of cells per motility state, we computed the Shannon entropy of each of the three age groups, which showed an increased cell-to-cell variation in young and post-adolescent/adults and a minimization of cellular heterogeneity for older adults (**Figure 4C**).

Together, these results demonstrate that age-associated cell motility states can be defined based on spatial and the activity patterns of cell movements, with the redistribution among these states partially explaining the cell-to-cell variations and heterogeneity observed as a function of age.

## DISCUSSION

Single-cell motility parameters are routinely reported as bulk values such as average displacements, speed, and persistence. Taking advantage of the single-cell nature of our motility measurements, we quantified changes in cell motility patterns not fully appreciated from standard bulk analysis (**Figure 1A-F**). Previously, Kimmel *et al*. employed single-cell analysis to show that mouse embryonic fibroblasts and myoblasts display heterogeneous phenotypic states across multiple scales with the potential for state-transitions in motility^18,20^. Taking a similar approach, we pooled the individual trajectories of all cells for all ages and identified eight spatial clusters (**Figure 2A-B**, **Supplementary Figure 4A**) and four activity clusters (**Figure 3C-E**) that classified groups of cells having similar motility phenotypes. Combining the spatial and activity patterns of single cells, we defined age-dependent motility states (**Figure 4A-B**). This approach highlights the immense amount of information that can be extracted at the single-cell level^5,19,28^, and demonstrates that dermal fibroblasts derived from healthy donors comprise a mixture of motile and non-motile cells, regardless of age (young, post-adolescent/adults or older adults).

This is an important finding since it provides a novel outlook of ageing at the cellular level, and a potential mechanism of how populations of cells encode and manifest age-dependent phenotypes. For instance, our data suggests that this decrease in overall motility with age is partly encoded by the weighted abundance of cells classified within spatially defined patterns of movement (**Supplementary Figure 5A-B**), with cells from young donors possessing a consistent movement pattern with periodic bursts of large displacements (above 1SD relative to baseline). These findings confirm our working hypothesis that the decrease in global cell motility with age is not due to the decrease in motility of all cells, but rather from the re-distribution of cells among motility states.

Assessing cell heterogeneity is critical to our understanding of emergent phenotypes in the context of health and disease^29–32^. Taking advantage of modern single-cell analysis approaches, we show that cell-to-cell variations of motility is a defining feature of ageing cells, with a minimization of cell heterogeneity for older adults (**Figure 4C**). Furthermore, developing a portrait of ageing at the single-cell level allows the investigation of novel questions regarding possible age-dependent phenotypic transitions. For instance, we wondered whether we could use the likely progression order among spatial clusters to identify age-associated motility tendencies (see Materials and Methods), such as whether the decrease in persistence of cells preluded the decrease in cellular displacement observed with increasing age. To address this, we computed the magnitude of the cross-correlation coefficients among each clusters (P1-P8) (**Supplementary Figure 4B**), and the strength of the correlation based on the abundances of each spatial cluster with age (**Supplementary Figure 4C-D**). Together, these correlation trends defined the likely transition order among clusters with age, and provides insights into the order of changes in persistence and displacements. The data suggests that cells tend to decrease their displacements before losing their ability to move in a persistent manner with increasing age (**Supplementary Figure 4E**).

In summary, we have demonstrated that using singe-cell approaches can lead to the identification of emergent patterns of cell motility as a function of age. This highlights the notion that single-cell approaches offer new insights into ageing at the cellular level. We anticipate that the increased implementation of modern single-cell analyses will lead to a more comprehensive understanding of ageing, and the ability to identify cellular states and phenotypic patterns that can be used as proxies of ageing in the context of health and disease. While these findings improve our present understanding of cellular determinants of ageing with regard to motility patterns, and its utility to serve as robust biomarkers of ageing, it remains unclear whether and how cells transition across motility states as a function of increasing age. This diversity in cellular phenotypes with age is likely linked to underlying molecular programs, cellular subtypes and cell cycle states that together influence the motility patterns of cells^5,33^. We anticipate that future work is needed to address this, which will require the use of cells derived from large cohorts of healthy donors (cross-sectional and longitudinal) imaged for long periods of time (order of days), coupled with single-cell molecular assessments.

## Supporting information

Supplementary dataset of xy coordinates for cells

## Acknowledgements

We acknowledge the financial support for this work from the National Institutes of Health; grant numbers U01AG060903 (DW, JW, JMP, PW), U54CA143868 (DW), R01CA174388 (DW), P30AG021334 Johns Hopkins Older Americans Independence Center (JW).

## Author contributions

JMP and DW conceived study design and analysis; JMP, MPP, JD, SMG, WW and KT performed experiments and tracked cells; JMP, NZ, and DW conceived analysis and analyzed data; JMP, DW, NZ, JW, and HJ interpreted results; DW, JW and PW supervised study; JMP and DW wrote manuscript; all authors contributed in reviewing and editing the manuscript.

## Code availability

Detailed descriptions of our approach and code utilized is either provided in the supplementary documentation or is already available through other published literature.

## Data availability

The authors declare that all data supporting the findings in this study are available within the paper and its Supplementary documents/information.

## MATERIALS AND METHODS

### Cell culture

A panel of twelve early-passage, primary dermal fibroblasts ranging in age from 2 to 92 years old (GM00969, GM05565, GM00038, GM00323, GM06111, AG04054, AG11796, AG08904, AG09162, AG12940, AG09558, AG09602), were obtained from Coriell Biobank cell repository (Camden, NY, USA), from collections comprising the Baltimore Longitudinal Study of Aging (BLSA) and the NIGMS apparently healthy controls. Cells were cultured in high-glucose (4.5 mg/ml) DMEM (Gibco), supplemented with 15% (vol/vol) fetal bovine serum (Hyclone), and 1% (vol/vol) penicillin-streptomycin (Gibco). Cell cultures were passaged every three to four days or when flasks were at ~80% confluence, for a maximum of five passages used for motility experiments. Data for cell trajectories for 860 cells across all ages can be found in Supplementary table S1.

### Quantification of cell motility

Fibroblasts were seeded at low density (~2000 cells/ml) onto type-1 collagen-coated (50μg/ml) substrates in 24-well plates and allowed to adhere overnight. Once cells attached, the plate was mounted unto a Nikon TE2000 microscope equipped with a motorized stage with X-Y-Z controls (Prior scientific) and environment control to maintain physiological conditions of temperature (37°C), Carbon dioxide (5%) and humidity (Pathology Devices). Phase-contrast images were recorded every 3 min for 16h using a Cascade 1K CCD camera (Roper Scientific) with a low magnification 10x Plan Fluor objective (numerical aperture, 0.3; Nikon). Cell motility parameters were determined from x-y coordinates obtained from the cell-centroid tracking of individual cells (MetaMorph). Cells undergoing division, moved out of frame, or had long contact times with other cells were omitted from analysis. Only cells having ten continuous hours of trackable movements that did not meet the above exclusion criteria were used for the final analysis. X-Y coordinates were then exported for analysis in Matlab for the quantification of spatial^24^ and activity parameters described below.

To quantify the spatio-temporal patterns of motility, we analyzed bulk cell movements based on trajectories using the Anisotropic Persistent Random Walk model (APRW)^24^. From this analysis we generated the parameters that describe the movements of the cells, (MSD60, MSD6, SP60, SP6, Pp, Pnp, Dp, Dnp, Dtot, ϕ), together with the mean-squared displacements, the auto-correlation function of velocities and the angular velocity magnitudes (see Glossary for definitions), which were computed and fitted based on the following equations:

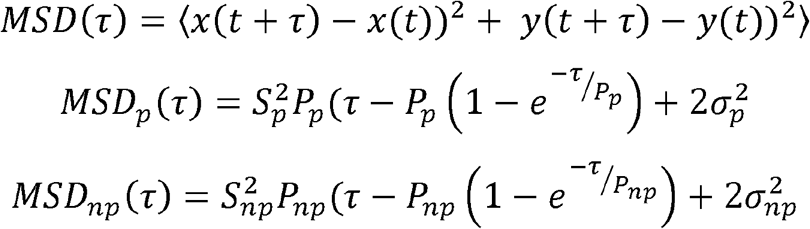

where S is the cell speed, *P* is the persistence time, 2*σ*^2^ is the noise (error) in the position of the cell, τ=nΔt and n=1,2, … N_min_−1, Δt is the size of the time step

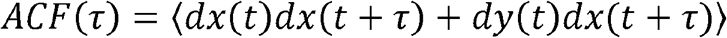

where

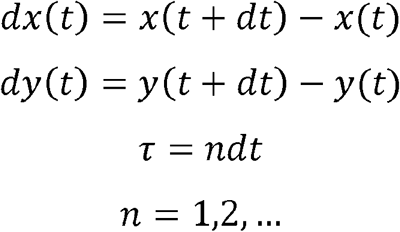

### Defining spatial motility clusters

Cell motility parameters describing the cells’ displacements, speeds, persistence times, diffusivities and the spatial persistence/anisotropy were computed using the anisotropic persistence random walk model (APRW), described previously by Wu et al^4,5^. For bulk motility analysis, ten motility parameters (MSD60, MSD6, SP60, SP6, Pp, Pnp, Dp, Dnp, Dtot, ϕ) were computed for each cell (see Glossary). For bulk analysis, parameters were averaged across all cells per age and the magnitude of the Pearson correlation coefficient was determined (**Figure 1F** and **Supplementary Figure 1**). For single-cell analyses, the distributions of motility parameters were log normalized to generate a normal distribution per parameter (**Supplementary Figure 2**). This resulted in the reduction of motility parameters from ten to eight (MSD60, MSD6, Pp, Pnp, Dp, Dnp, Dtot, ϕ), since the normalized MSD6 was roughly equal to SP6, and MSD60 equal to SP60, (i.e. 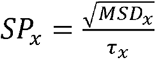, where τ=time lag). Following the normalization, the eight motility parameters were used to define the spatial clusters (Pn). Spatial clusters were defined based on the abundance of cells having similar magnitudes of the eight features defined in the APRW model. Specifically, the motility features were computed for each cell and compiled for all cells across all ages. Importing this data into Matlab, we performed unsupervised hierarchical clustering analysis based on the Cityblock distances along the axis of maximum variation (ward linkages). This clustering analysis resulted in the stratification of cells into eight clusters (P1-P8).

### Determining the likely progression order

To determine the likely progression order with age, we computed the magnitude of the correlation for the abundance of cells per cluster with age, and the cross correlation among clusters. Once the correlation coefficients were determined, the correlations of cell abundances with age were ranked to determine the overall progression order. In addition to determine the linkages (length—strength of cross-correlation) of clusters we also ranked the correlations among clusters to determine the cluster-to-cluster proximity. Once both sets of correlations were compiled, a network were constructed such that the size of the nodes were scaled based on the number of cells in each cluster, and the cluster-cluster proximity denoted the strength of the cross correlation. The magnitudes of the Pearson correlation coefficients were computed in Matlab, (R=corrcoef(A), where ‘R’ is the Pearson correlation coefficient and ‘A’ is the data matrix of cell abundances.

### Quantifying the activity profiles per single cell

To determine the activity profiles of individual cells, the raw x-y trajectories for each cell were converted into 1-dimensional displacement trajectories (see **Figure 3A-B**). Here, the temporal displacement frequencies and the presence/absence of spikes (burst of movement) and trains (continuous bursts of movements) defines the activity space. We computed the activity based on the displacements according to this equation:

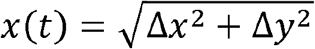

where, Δ*x* = *x*(*t*) − *x*(*t* − 1) and Δ*y* = *y*(*t*) − *y*(*t* − 1), are the changes in the vector components of the cell movements in Cartesian coordinates at time *t*, Δ*t* is the time step between different measurements of cell positions.

Once this was computed for each cell, the magnitudes of the displacements were z-score normalized per unit time so that each cell was normalized to its own baseline movements, and comparable on the same numerical scale.

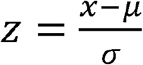

where, z is the z-score, x is the magnitude of the variable, μ and σ denote the mean values and the standard deviation respectively.

The activity was computed for all 860 single cell across all ages, then we performed unsupervised hierarchical clustering analysis to delineate groupings of cells having similar activity profiles. This was done in Matlab based on the Cityblock distances along the axis of maximum variation (ward) which yielded five clusters that we later interrogated to identify cluster-dependent motility patterns.

To calculate the binarized activity, the continuous activity profile was transformed into a binary matrix of 1’s and 0’s denoting trains and lags. Trains denote bursts of movements one standard deviation above the baseline movement, and lags denotes time steps at baseline and below the threshold. Distribution of trains and lags were computed by compiling the series of 1’s and 0’s within each temporal activity pattern for all cells and compiles per activity cluster, respectively.

### Computing cellular heterogeneity

To quantify the heterogeneity among cells (both for spatial and activity clusters) we utilized the Shannon entropy. Here the entropy *S* was calculated as:

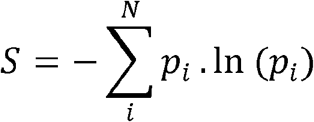

where *p_i_* corresponds to the fractional abundance of cells within the particular cluster, for each age group N (young, middle age and older adults). The entropy was determined on both a per age and age-groups, based on the abundance of cells per cluster (i.e. both spatial and activity). Entropy per age group was computed by taking the average of entropies per age (young-A02, A03, A09, A11, A16; post-adolescent/adults-A29, A35, A45, A55; older adults-A65, A85, A92).

### Quantifying enrichments and depletions of motility clusters and states

To determine whether the clusters defined above (spatial and activity) were significantly depleted or enriched as a function of age or age groups, we utilize a randomization strategy. For each cluster ‘i’, we obtain 2 p-values per age corresponding to null hypotheses of under-representation (depletion) and over-representation (enrichment), respectively. Specifically, for each cluster ‘i’ and for each age or age-group ‘k’, we compare the observed frequency of cells with a distribution of expected frequencies of cells of age k, built by from N samples (50,000 permutations) of the size of cluster ‘i’.

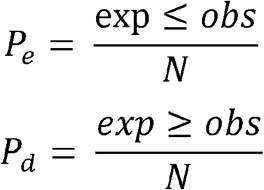

where, ‘exp’ denotes the simulated abundance based on the null hypothesis, and ‘obs’ denoted the observed abundance of cells per cluster/state. P_e_ and P_d_ denote the p-values for the enrichment and depletion, respectively.

### Statistics and rigor

All experiments were conducted with in-plate technical controls in triplicates. The specific number of cells used in analysis are denoted in the main figures for each section. Correlation analysis was conducted using the Pearson correlation coefficients and statistical significance were assessed using either t-tests or one-way ANOVA. To compute the significant enrichments and depletions within each of the 40 defined motility states, we employed a randomization enrichment test for 50,000 permutations and compared to observed frequencies. Significance was determined based on the magnitude of the p-values (* p<0.05, ** p<0.01, *** p<0.001).

## GLOSSARY

MSD6: Mean-squared displacements at time lag equal to 6 minutes
MSD60: Mean-squared displacement at time lag equal to 60 minutes
Pp: Persistence time along the primary axis of migration
Pnp: Persistence time along the secondary axis of migration
PRW: Persistence random walk
APRW: Anisotropic persistence random walk
Mean-squared displacement (MSD): the average displacement of a cell per unit time lag (*dt*), which is a multiple of the time step. The MSD is a common measure of random movements of cells.
Autocorrelation function of velocity (ACF): is a measure of the correlation in cell velocities per unit time lag (*dt*). Higher values of the ACF typically indicate that cells are more persistent in their movement.
Angular velocity magnitude: is a measure of the magnitude of average cell velocities evaluated at different orientations after re-alignment along the primary axis of migration. This reveals the degree of anisotropy of cell velocities. If the velocity profile is isotropic (circular), as is typical in cases of persistent random walks, the average magnitude of the velocity is equally likely in all directions. However, if the velocity is anisotropic, as seen for cells from young donors, the average velocity along the primary axis is significantly higher that along other axes of movements.
Spatial clusters (Pn): Motility clusters defined based on the eight motility parameters defined using the APRW model
Activity clusters (ACn): Motility clusters defined by the 1D displacement profiles of single cells that define the temporal motility patterns
Shannon entropy (S): A surrogate measurement of cellular heterogeneity based on the number of states and the abundances of cells within each state
SP6: Average speed computed at a 6-min time lag

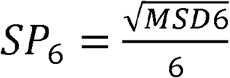
SP60: Average speed computed at a 60-min time lag

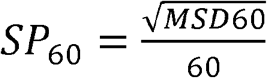
Dp: Diffusivity along the primary axis of migration (motility coefficient)

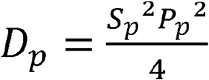
Dnp: Diffusivity along the secondary axis of migration (motility coefficient)

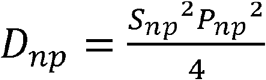
Dtot: Total diffusivity, equal to the sum of diffusivities along the primary and secondary axes of migration

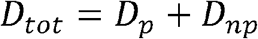
Psi (ϕ): Anisotropy (spatial persistence), equal to the ratio of the diffusivities in the primary and secondary axes of migration

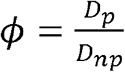
Trains: length of time steps within the normalized activity profile per cells having a value greater than or equal to one standard deviation above the baseline.
Lags: length of time steps within the normalized activity profile per cells having a value less than one standard deviation above the baseline.

## SUPPLEMENTARY FIGURE CAPTIONS

**Supplementary Figure 1.**
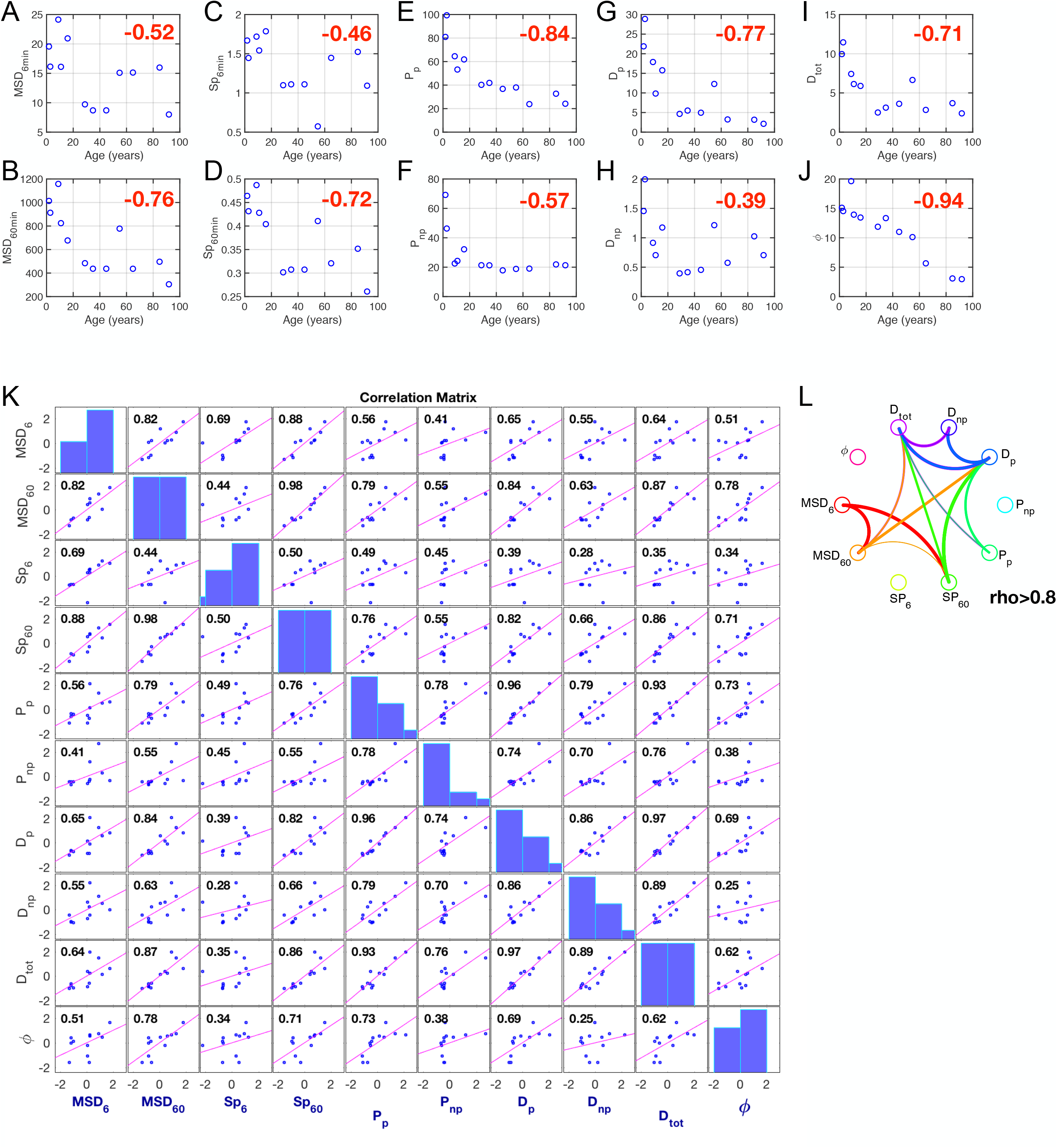
Cross correlation analysis of motility parameters with age. **A-J.** Pearson correlation analysis of global cell-motility parameters with increasing age; MSD6 (**A**), MSD60 (**B**), SP6 (**C**), SP60 (**D**), Pp (**E**), Pnp (**F**), Dp (**G**), Dnp (**H**), Dtot (**I**), ϕ (**J**). **K.** Cross correlation analysis across parameters with age; each dot represents a donor, with Pearson correlation coefficients denoted in the upper left of the figure. **L**. Circus plot showing the magnitudes of the correlation coefficients among parameters, connected nodes denote cross correlations above 0.8, with the thickness of the lines showing the scaled magnitude.

**Supplementary Figure 2.**
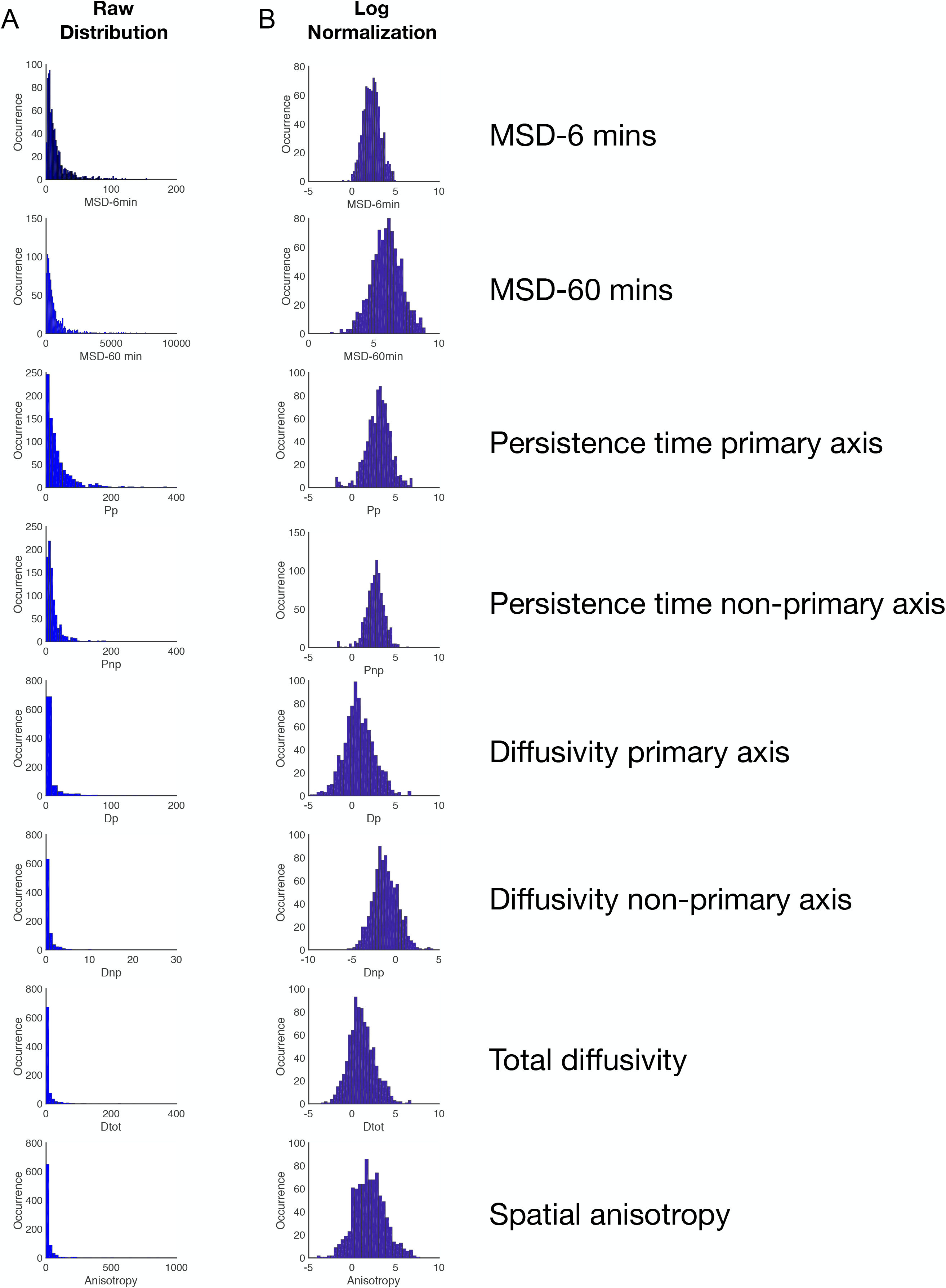
Log normalization of cellular motility parameter distributions. **A**. Raw distributions of eight motility parameters. **B**. Log normalization of the corresponding distributions for motility parameters.

**Supplementary Figure 3.**
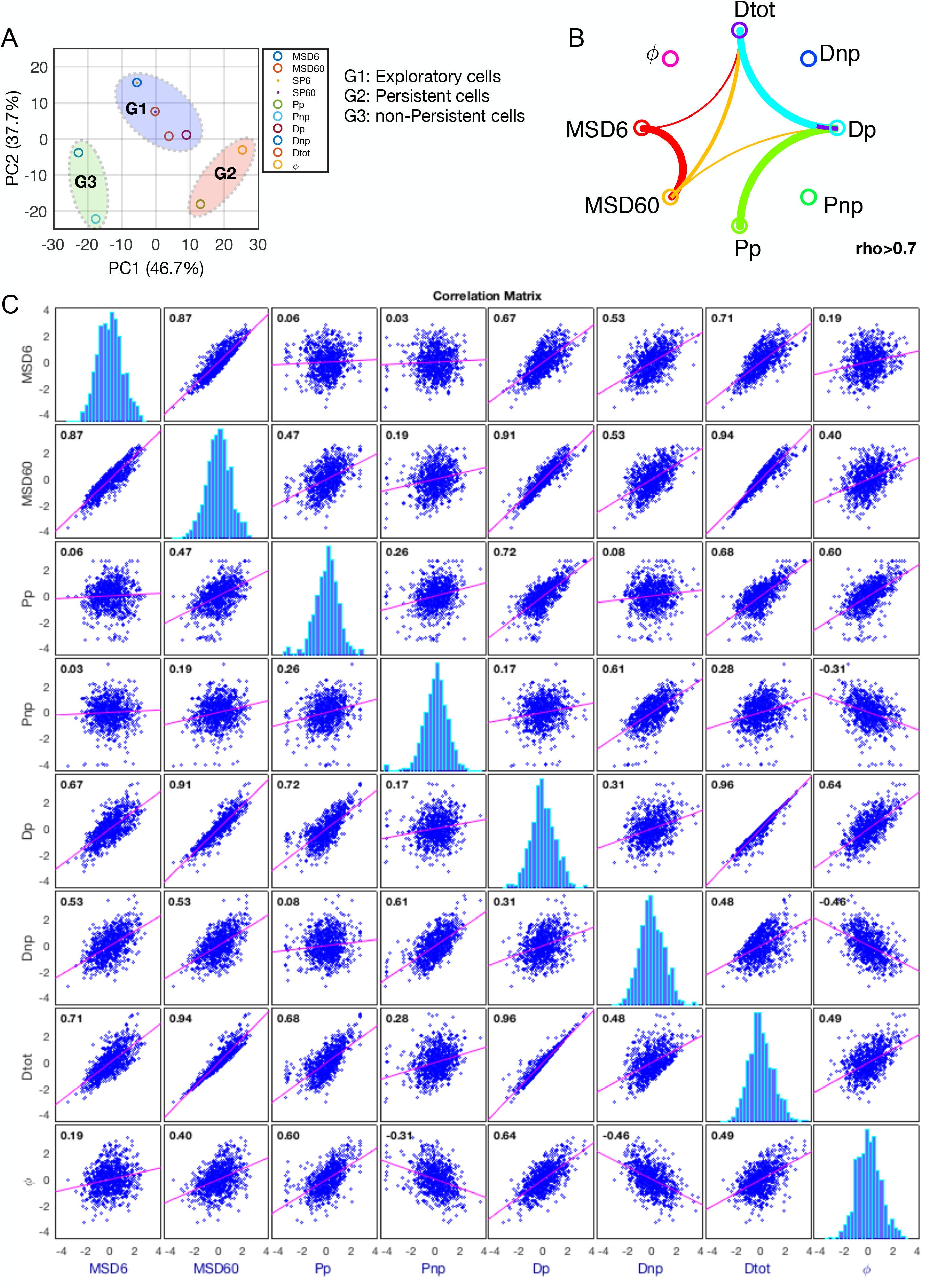
Correlation analysis of motility parameters at the single-cell level. **A.** Principal component analysis (PCA) of the eight motility parameters showing the delineation into three groups, G1-high displacement cells, G2-persistent, and G3-non-persistent. **B**. Circus plot showing the cross correlation among the eight motility parameters at single cell resolution. Lines connecting nodes denote correlation coefficients greater than 0.7. **C.** Cross correlation analysis among eight motility parameters across all ages, each dot represents a single cell, with the linear regression line shown in magenta, and the magnitude of the Pearson correlation coefficient displayed in the upper left corner of the plots.

**Supplementary Figure 4.**
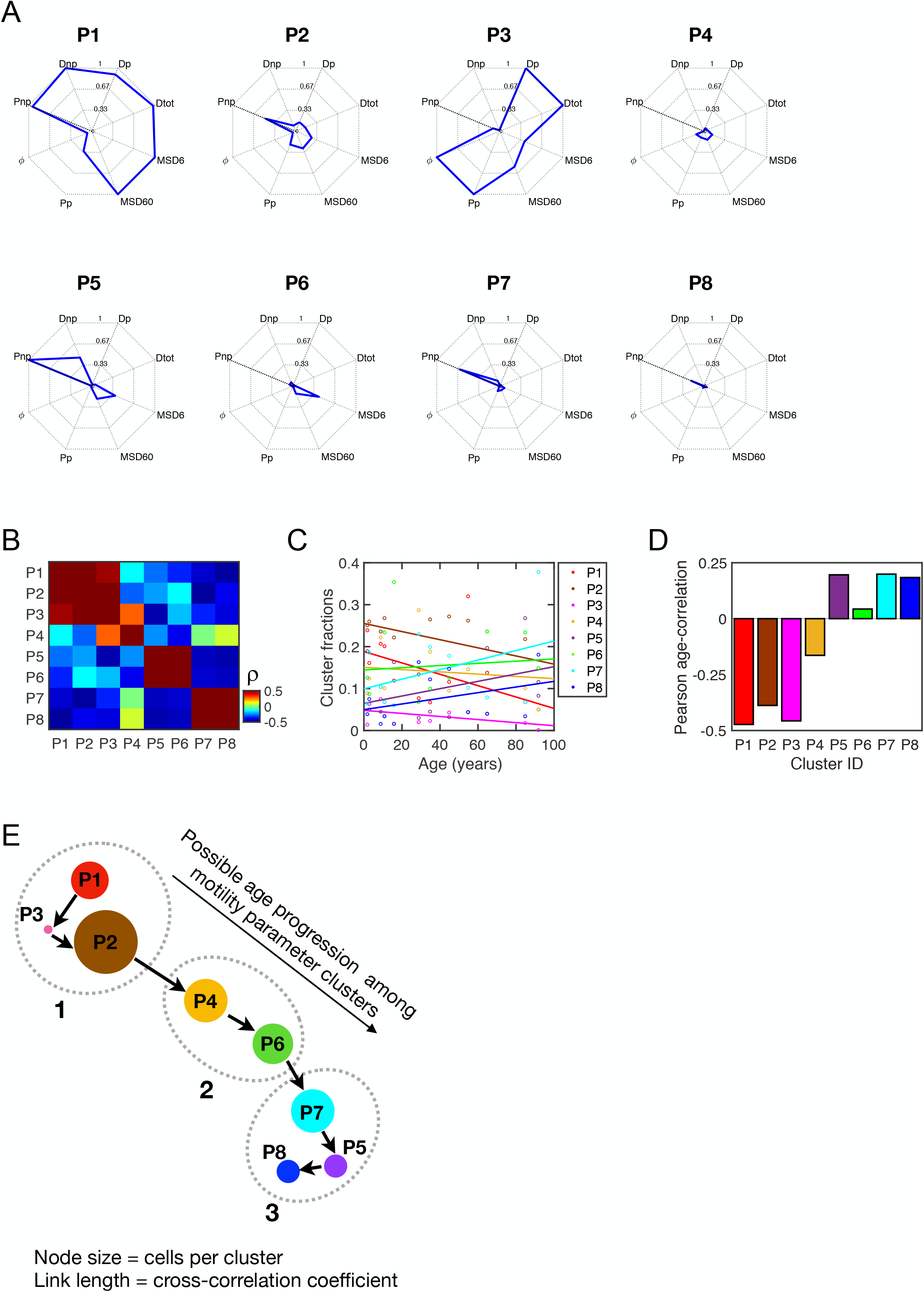
Single cell spatial motility clusters are defined by similarities in the magnitude of motility clusters. **A.** Radar plots showing the normalized magnitudes (z-score) for the eight motility parameters defining the eight distinct motility clusters. **B**. Heatmap showing the magnitude of the age correlation among eight clusters. Color scale from blue to maroon denoting low to high, with a range of Pearson correlation coefficients from −0.5 to 0.5. **C**. Pearson correlation trends showing the relationship of cell fraction within each color-coded cluster with increasing age. **D**. Bar plots showing the magnitude of the age correlations for each cluster shown in (B). **E**. Proposed progression order of cells within each cluster with age. Since aging is a continuum, the order was determined based on the magnitude of the correlation for each of the 8 clusters with increasing age, and the associating among clusters (how close or far from each other) was determined based on the cross correlation among clusters across all ages. Results indicate that the likely progression order is P1, P3, P2, P4, P6, P7, P5, P8. **F**. Bar plot showing the proportion of cells being classified in each spatial and activity cluster.

**Supplementary Figure 5.**
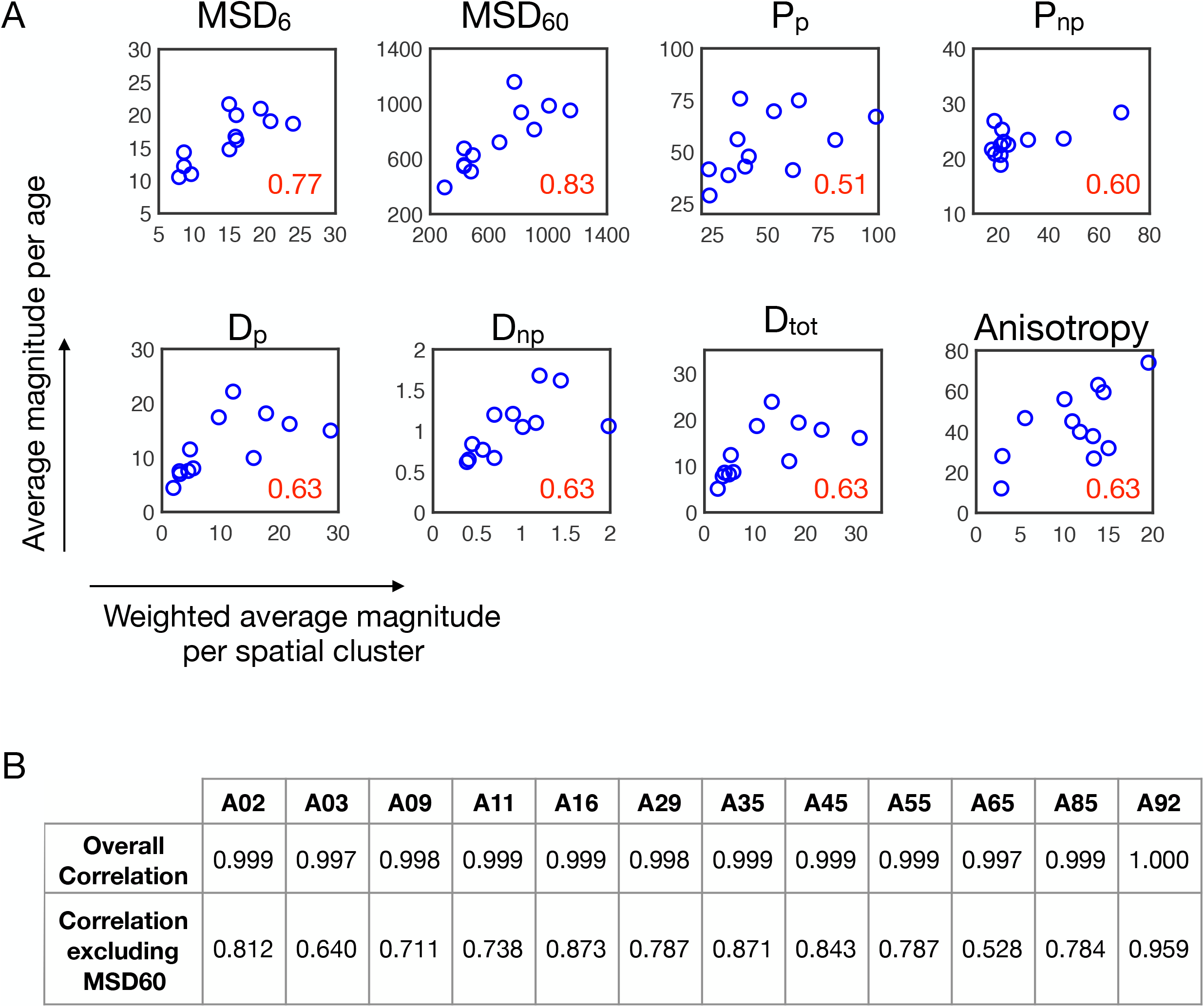
Age-associated motility parameters scale as weighted averages per spatial cluster. **A.** Scatter plots showing the correlation between the weighted average per spatial cluster and the average magnitude of the parameter cluster. Pearson correlation coefficient is denoted in red in the bottom right side of plots. The weighted average was calculated as the sum of the fractions times the average magnitude per cluster. **B**. Table showing the correlation of the sum of the weighted averages per cluster with all average motility parameters as a function of age.

**Supplementary Figure 6.**
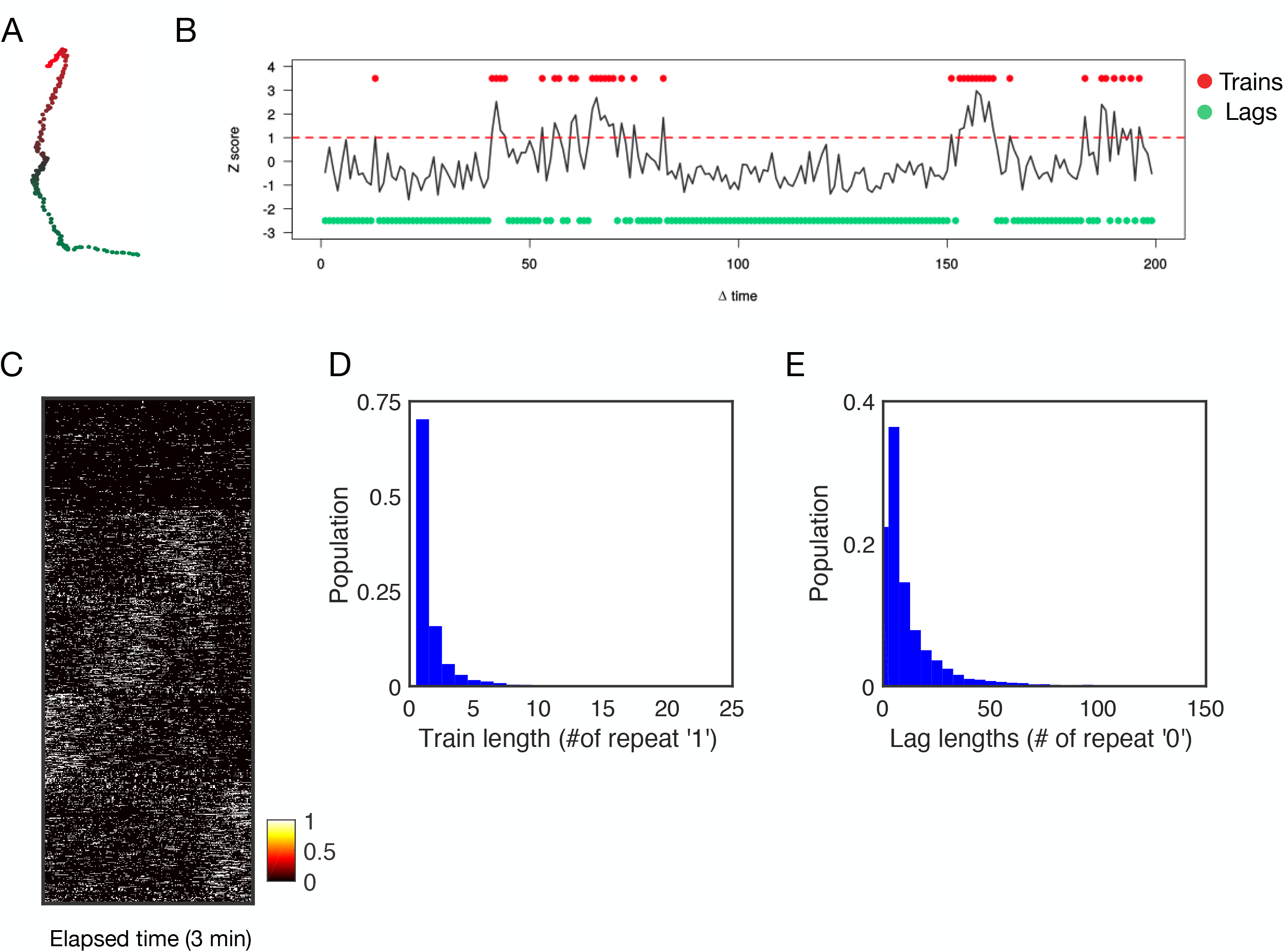
Cellular activity based on point-process analysis. **A**. x-y trajectory of a single cell. **B.** 1-Dimensional displacement profile for cells shown in (A), highlighting how we convert the raw activity profile is transformed into a binary profile based on a threshold of 1 standard deviation above the baseline. Red dashed line denotes 1SD, with red circles denoting trains and green circles denoting lags. **C**. Heat map showing the binary activity profiles per cell, ordered based on 4 activity groups show in Figure 3C. **D-E**. Distribution of train lengths (D) and lag lengths (E) across all cells for all ages.

**Supplementary Figure 7.**
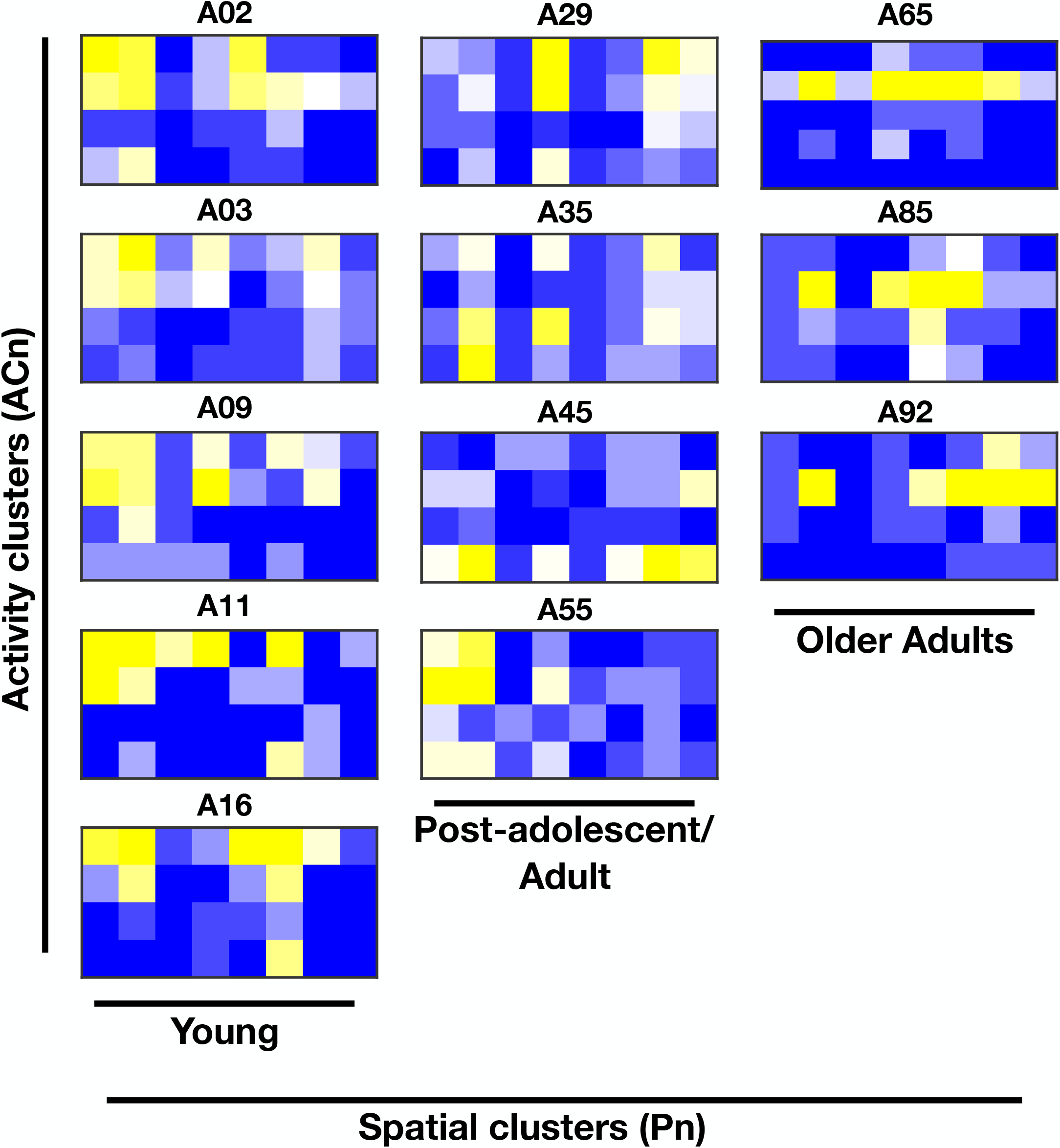
Association among spatial and activity clusters per age. Heatmaps showing the frequencies of cells per motility state for all ages. Each abundance is normalized based on the maximum value of frequencies per age.

**Supplementary Table 1.** Name and age for samples used in this study

**Supplementary Table 2.** Enrichments and depletions for each spatial cluster per age

**Supplementary Table 3.** Enrichments and depletions for each activity cluster per age

**Supplementary Table 4.** Enrichments and depletions for spatial clusters per activity cluster

**Supplementary Table 5**. Enrichments and depletions for spatial clusters per activity cluster for each age group

**Supplementary dataset 1.** Table of X-Y coordinates for all 860 single cells across all ages

## REFERENCES

1. Maynard, S., Fang, E. F., Scheibye-Knudsen, M., Croteau, D. L. & Bohr, V. A. DNA damage, DNA repair, aging, and neurodegeneration. Cold Spring Harb. Perspect. Med. (2015). doi:10.1101/cshperspect.a025130

2. López-Otín, C., Blasco, M. A., Partridge, L., Serrano, M. & Kroemer, G. The hallmarks of aging. Cell 153, (2013).

3. Phillip, J. M., Aifuwa, I., Walston, J. & Wirtz, D. The Mechanobiology of Aging. Annu. Rev. Biomed. Eng. 17, 113–141 (2015).

4. Valdes, A. M., Glass, D. & Spector, T. D. Omics technologies and the study of human ageing. Nature Reviews Genetics (2013). doi:10.1038/nrg3553

5. Mahmoudi, S. et al. Heterogeneity in old fibroblasts is linked to variability in reprogramming and wound healing. Nature (2019). doi:10.1038/s41586-019-1658-5

6. Belsky, D. W. et al. Quantification of biological aging in young adults. Proc. Natl. Acad. Sci. 112, E4104–E4110 (2015).

7. Alpert, A. et al. A clinically meaningful metric of immune age derived from high-dimensional longitudinal monitoring. Nat. Med. (2019). doi:10.1038/s41591-019-0381-y

8. Zhou, W. et al. Longitudinal multi-omics of host–microbe dynamics in prediabetes. Nature (2019). doi:10.1038/s41586-019-1236-x

9. Horvath, S. DNA methylation age of human tissues and cell types. Genome Biol. (2013). doi:10.1186/gb-2013-14-10-r115

10. Ahadi, S. et al. Personal aging markers and ageotypes revealed by deep longitudinal profiling. Nature Medicine (2020). doi:10.1038/s41591-019-0719-5

11. Fleischer, J. G. et al. Predicting age from the transcriptome of human dermal fibroblasts. Genome Biol. (2018). doi:10.1186/s13059-018-1599-6

12. Lehallier, B. et al. Undulating changes in human plasma proteome profiles across the lifespan. Nature Medicine (2019). doi:10.1038/s41591-019-0673-2

13. Bhuva, A. N. et al. Training for a First-Time Marathon Reverses Age-Related Aortic Stiffening. J. Am. Coll. Cardiol. (2020). doi:10.1016/j.jacc.2019.10.045

14. Epel, E. S. Can childhood adversity affect telomeres of the next generation? Possible mechanisms, implications, and next-generation research. American Journal of Psychiatry (2020). doi:10.1176/appi.ajp.2019.19111161

15. Rosero-Bixby, L. et al. Correlates of longitudinal leukocyte telomere length in the Costa Rican Longevity Study of Healthy Aging (CRELES): On the importance of DNA collection and storage procedures. PLoS One 14, e0223766 (2019).

16. Schüssler-Fiorenza Rose, S. M. et al. A longitudinal big data approach for precision health. Nat. Med. (2019). doi:10.1038/s41591-019-0414-6

17. Phillip, J. M. et al. Biophysical and biomolecular determination of cellular age in humans. Nat. Biomed. Eng. 1, 0093 (2017).

18. Kimmel, J. C., Chang, A. Y., Brack, A. S. & Marshall, W. F. Inferring cell state by quantitative motility analysis reveals a dynamic state system and broken detailed balance. PLoS Comput. Biol. (2018). doi:10.1371/journal.pcbi.1005927

19. Chen, X. et al. Single-cell analysis at the threshold. Nature Biotechnology (2016). doi:10.1038/nbt.3721

20. Kimmel, J. C., Hwang, A. B., Marshall, W. F. & Brack, A. S. Aging induces aberrant state transition kinetics in murine muscle stem cells. bioRxiv (2019). doi:10.1101/739185

21. Kim, D.-H., Cho, S. & Wirtz, D. Tight coupling between nucleus and cell migration through the perinuclear actin cap. J. Cell Sci. 127, 2528–2541 (2014).

22. Khatau, S. B. et al. The distinct roles of the nucleus and nucleus-cytoskeleton connections in three-dimensional cell migration. Sci. Rep. (2012). doi:10.1038/srep00488

23. Pienta, K. J. & Coffey, D. S. Characterization of the subtypes of cell motility in ageing human skin fibroblasts. Mech. Ageing Dev. (1990). doi:10.1016/0047-6374(90)90001-V

24. Wu, P. H., Giri, A. & Wirtz, D. Statistical analysis of cell migration in 3D using the anisotropic persistent random walk model. Nat. Protoc. (2015). doi:10.1038/nprot.2015.030

25. Wu, P.-H., Giri, A., Sun, S. X. & Wirtz, D. Three-dimensional cell migration does not follow a random walk. Proc. Natl. Acad. Sci. 111, 3949–3954 (2014).

26. Shannon, C. E. A Mathematical Theory of Communication. Bell Syst. Tech. J. (1948). doi:10.1002/j.1538-7305.1948.tb01338.x

27. Villegas, P., Di Santo, S., Burioni, R. & Muñoz, M. A. Time-series thresholding and the definition of avalanche size. Phys. Rev. E (2019). doi:10.1103/PhysRevE.100.012133

28. Zhu, C., Preissl, S. & Ren, B. Single-cell multimodal omics: the power of many. Nature Methods (2020). doi:10.1038/s41592-019-0691-5

29. Altschuler, S. J. & Wu, L. F. Cellular Heterogeneity: Do Differences Make a Difference? Cell 141, 559–563 (2010).

30. Wu, P.-H. et al. Evolution of cellular morpho-phenotypes in cancer metastasis. Sci. Rep. 5, 18437 (2016).

31. Huang, S. Non-genetic heterogeneity of cells in development: more than just noise. Development (2009). doi:10.1242/dev.035139

32. Wu, P.-H. et al. Single-cell morphology encodes metastatic potential. Sci. Adv. (2020). doi:10.1126/sciadv.aaw6938

33. Efremova, M. & Teichmann, S. A. Computational methods for single-cell omics across modalities. Nature Methods (2020). doi:10.1038/s41592-019-0692-4

